## Supplemental Text, Figures and Tables for "TLR5 participates in the TLR4 receptor complex and biases towards MyD88-dependent signaling in environmental lung injury"

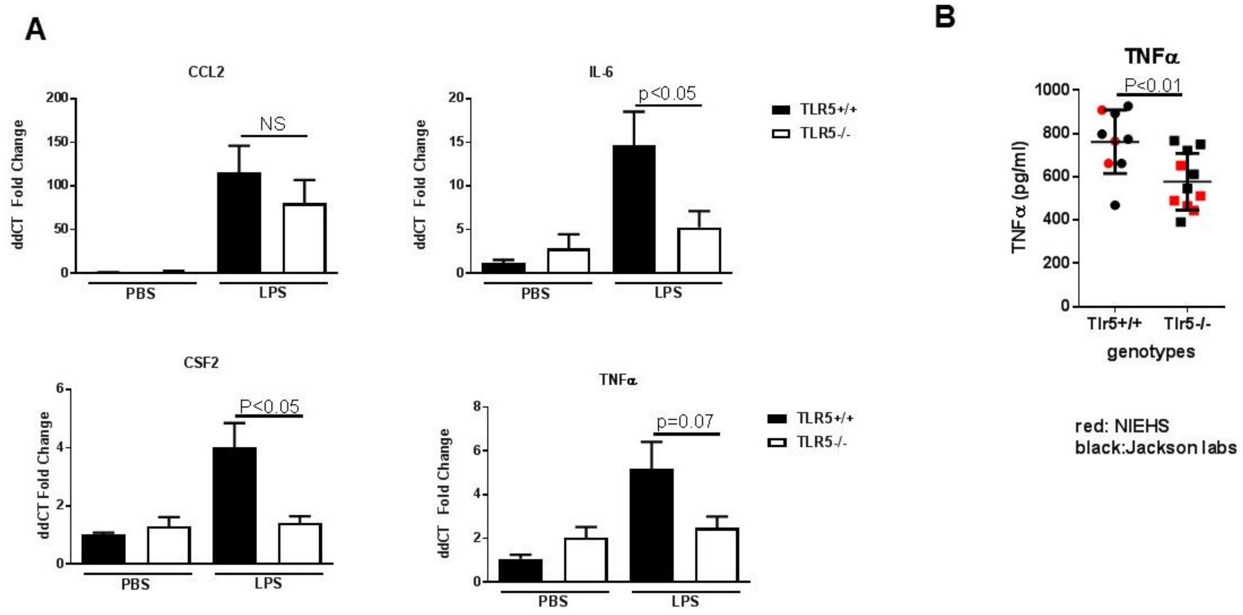


**Supplemental Figure 1. TLR5 deficiency ameliorates the inflammatory response to LPS exposure.** (**A**) Real time quantitative PCR analysis of inflammatory cytokines shows a significant decrease in *Tlr5*-deficient (TLR5$^{-/-}$) mice. N=5-8 mice per group, experiment repeated twice. Data are represented as mean ± s.e.m. and were analyzed by unpaired t test with Welch's correction. (**B**) Mouse provenance does not impact *Tlr5*-sufficient (Tlr5$^{+/+}$) and -deficient (Tlr5$^{-/-}$) differences in LPS response. NS=not significant

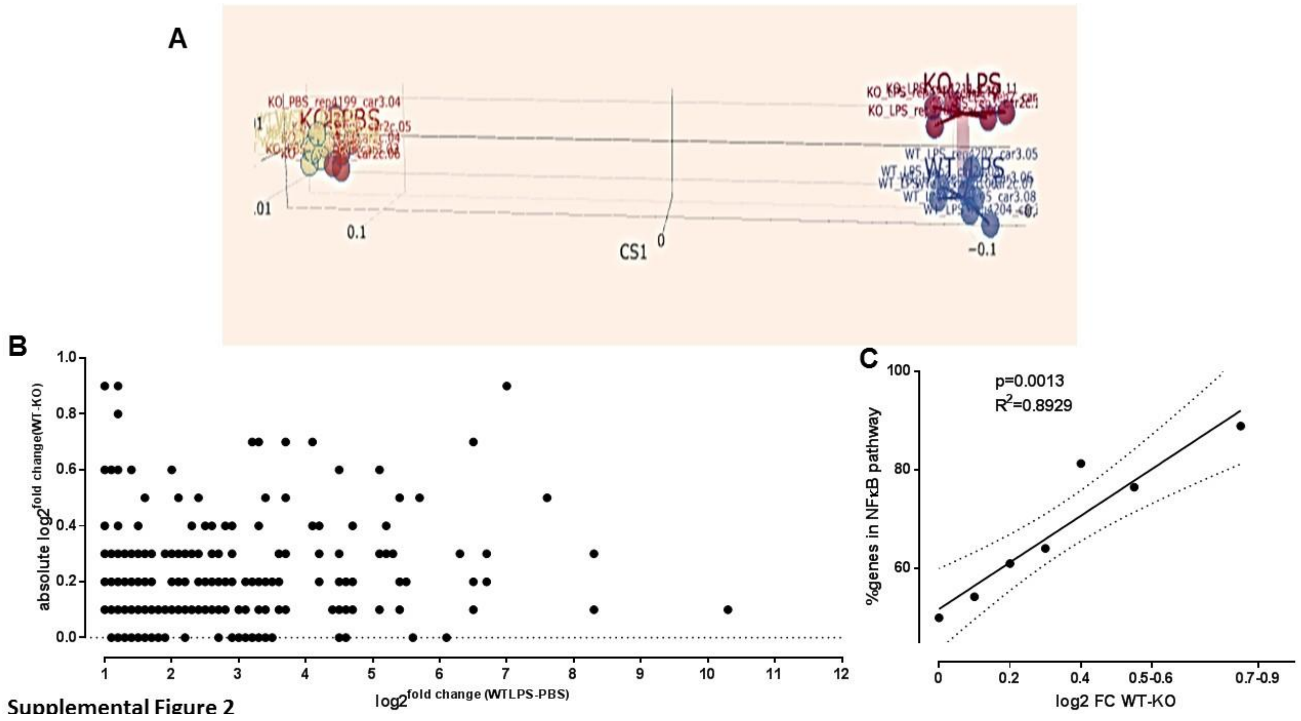


**Supplemental Figure 2. Immune gene expression profiling dependent on TLR5 after ultrapure LPS exposure.** mRNA from lungs of mice 3 hours after instilled ultrapure LPS exposure was analyzed using the NanoString© platform (www.nanostring.com) utilizing the Mouse Innate Immunity Panel Codeset (Ns_Mm_Myeloid_v2.0) and focusing on genes that were more than 2x upregulated after LPS exposure. N=6-7 mice per genotype and exposure. (**A**) Between Groups Analysis (BGA) clustering if gene expression after PBS and LPS exposure. BGA is a clustering method similar to principal components analysis (PCA), except that it prioritizes the separation of sample group centroids, as opposed to individual samples as with standard PCA. There is no clear separation of gene expression clusters after PBS exposure (left) but a clear separation after LPS exposure between *Tlr5*-deficient (KO, purple circles, right top) and *Tlr5*-sufficient (WT, blue circles, right bottom) (**B**) Distribution of TLR5 effects (y-axis, absolute Log2 values of the fold-difference between wildtype (WT) and Tlr5 deficient (KO) mice) plotted against the LPS-induced gene upregulation in WT mice after LPS exposure (x-axis, Log2 values of fold change in gene expression between PBS and LPS). (**C**) Proportion of the genes that are downstream of the NFκB pathway is inversely proportional to the magnitude of the effect of Tlr5. Among the genes that are 60% to 90% upregulated in WT mice vs. KO mice (log2 fold-change (FC) WT-KO=0.7-0.9), almost 90% are downstream of NFκB. This proportion falls to 50% among the genes that are not different between genotypes (log2FCWT-KO=0)

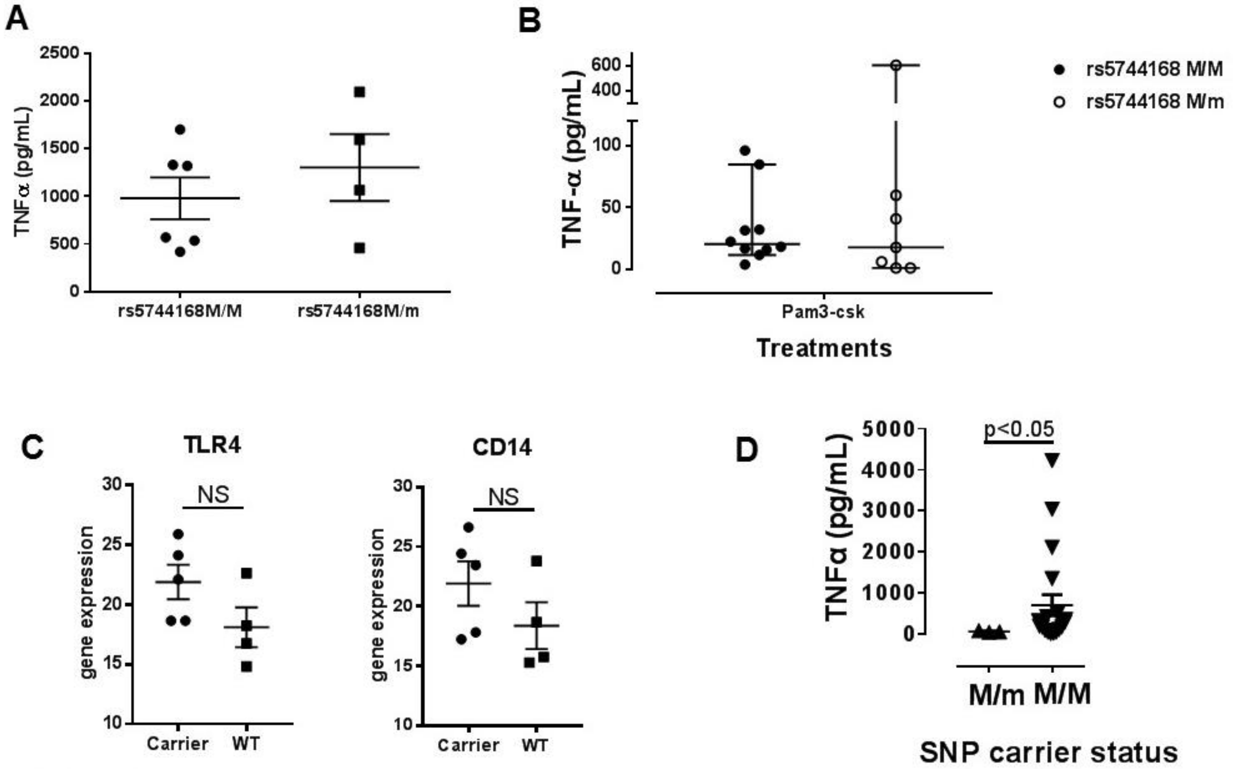


**Supplemental Figure 3. TNF secretion from monocyte-derived macrophages from human subjects depending on TLR5 status** (**A)** TNF-α secretion from whole blood cells from normal (rs5744168 M/M) or minor-allele-carriers (rs5744168 M/m) for TLR5 single nucleotide polymorphism after 6 hours ultrapure LPS exposure (10 ng/mL). Data are represented as mean mean ± s.e.m. for n = 6 individuals for rs5744168 M/M and n = 4 individuals for rs5744168 M/m. (**B)** TNF-α secretion by monocyte-derived macrophages from human volunteers either homozygote for the major allele (rs5744168 M/M) or carriers of the minor allele (rs5744168 M/m) for the TLR5 single nucleotide polymorphism rs5744168. Cells were exposed to 1 µg/ml Pam3CSK4 for 24 hours and TNF-α levels were analyzed by Duoset ELISA kit. Data are represented as median ± 95% CI and analyzed by unpaired t test with Welch's correction. N=7-10 individual subjects. (**C**) TLR4 and CD14 expression are not significantly different between monocyte-derived macrophages from human volunteers either homozygote for the major allele (rs5744168 M/M) or carriers of the minor allele (rs5744168 M/m) for the TLR5 single nucleotide polymorphism rs5744168. (**D**) TNF-α secretion by alveolar macrophages after ozone exposure: Deficiency is associated with decreased expression (analyzed by Mann-Whitney test).

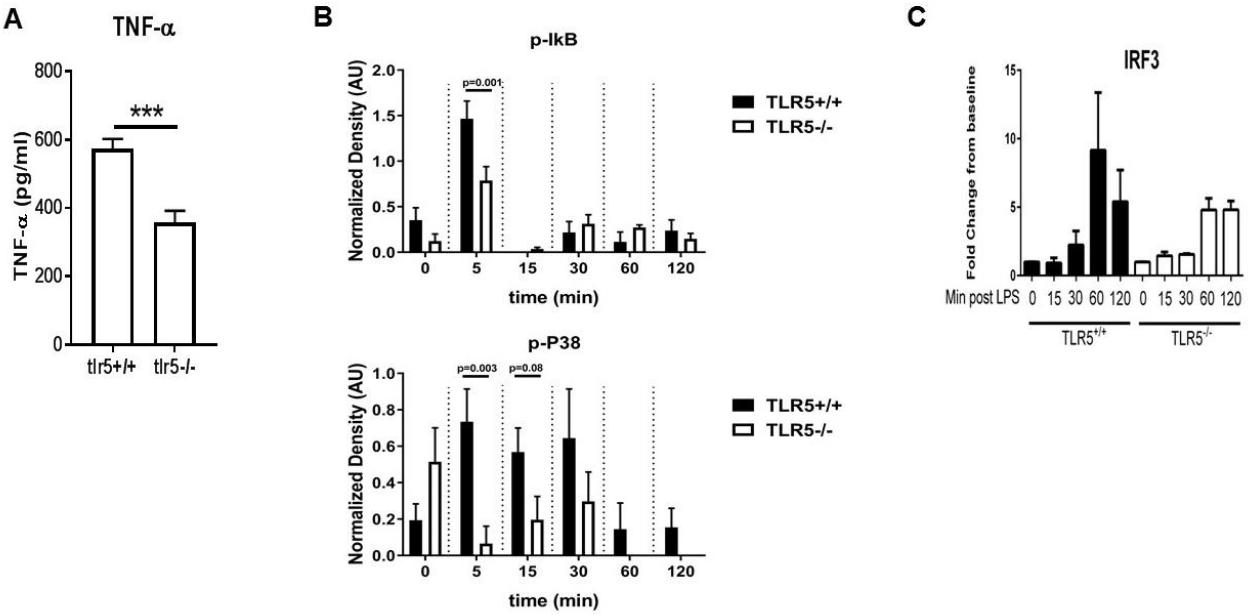


**Supplemental Figure 4.** (**A**) TNFα expression after exposure of *Tlr5*-sufficient (Tlr5$^{+/+}$) and -deficient (Tlr5$^{-/-}$) BMDM to monophosphoryl lipid A (MPLA). (**B**) Quantification of densitometric analysis for p-IkB and p-P38 of 3 separate blots similar to **(Fig. 3A).** (**C** ) Nuclear abundance of IRF3 in BMDM cells isolated from TLR5$^{-/-}$and TLR5$^{+/+}$ mice and matured ex vivo in the presence of 10 ng/mL murine mCSF. Cells were treated for 0,15,30,60, 120 minutes with ultrapure LPS (10 ng/mL), lysed and nuclear fractions were isolated. Isolated nuclear proteins were blotted with antibody against IRF3 (Cell signaling). Data are represented as mean ± s.e.m. and were analyzed by repeated unpaired t test with Holm-Sidak correction.

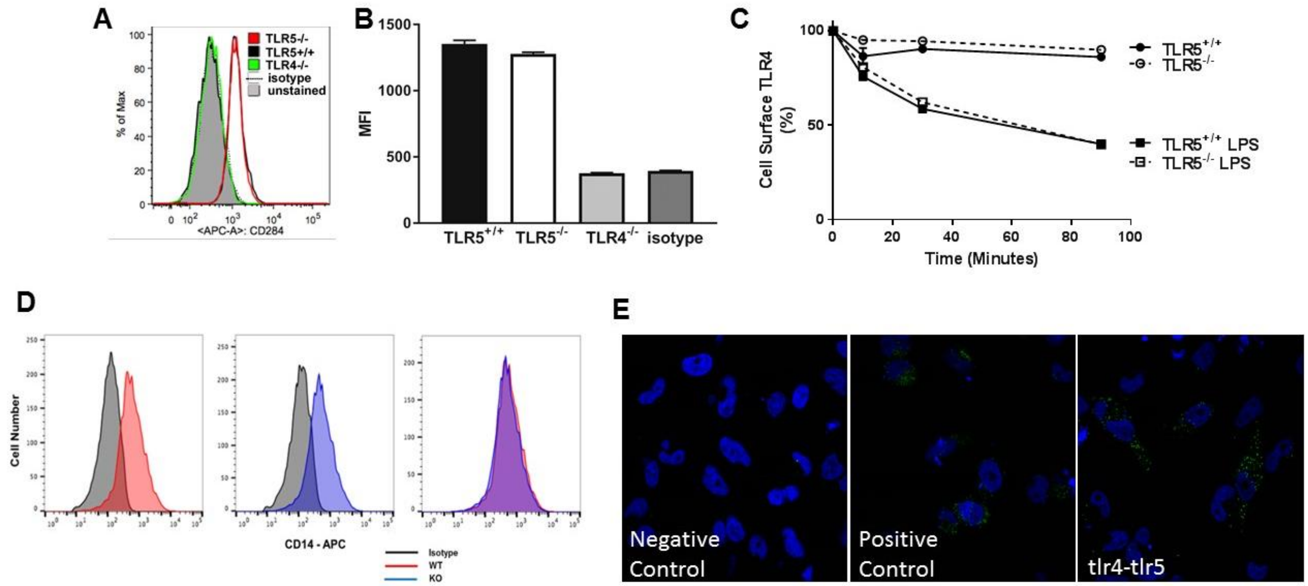


**Supplemental Figure 5. Cell surface expression of TLR4 and its co-localization with TLR5** (**A**) Cell surface TLR4 expression is not dependent on TLR5. Cells were isolated from *Tlr5*$^{-/-}$, *Tlr5*$^{+/+}$ and *Tlr4*$^{-/-}$ mice and matured *ex vivo* in the presence of 10 ng/mL murine MCSF. Cells were stained with either CD284 TLR4 antibody (Biolegend) or Isotype control. (**B**) Median fluorescence intensity analysis from the experiment present in panel a. Data are presented as mean ± s.e.m. for n = 9 mice per genotype. (**C**) TLR4 cell surface expression after 100 ng/mL ultrapure LPS exposure for up to 90 minutes in *Tlr5*-deficient (TLR5$^{-/-}$) or *Tlr5*-competent (TLR5$^{+/+}$) BMDM. (**D)** Cell surface CD14 expression is not dependent on TLR5. Cells were isolated from *Tlr5*$^{-/-}$ and *Tlr5*$^{+/+}$ mice and matured *ex vivo* in the presence of 10 ng/mL murine MCSF. Cells were stained with either CD284 TLR4 antibody (Biolegend) or Isotype control. (**E)** Proximity ligation assay for TLR4 and TLR5 in HeLa cells overexpressing TLR4-FLAG M2or MyD88-V5 and TLR5-HA.

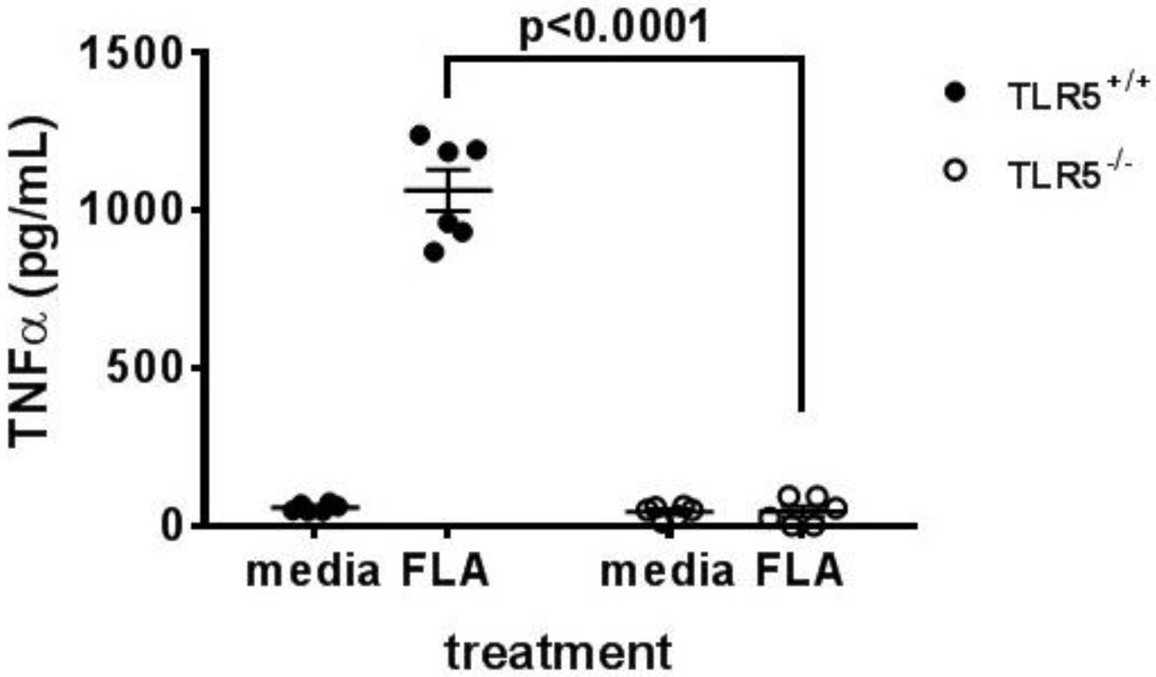


**Supplemental Figure 6. TLR5 deficient BMDM are not responsive to flagellin.** BMDM from either *Tlr5*-deficient (TLR5$^{-/-}$) or *Tlr5*-competent (TLR5$^{+/+}$) mice were exposed to 100 ng/ml ultrapure flagellin for 24 hours and TNF-α levels were analyzed by Duoset ELISA kit. Data are represented as mean ± standard deviation and analyzed by unpaired t test using the Holm-Sidak method. N=6 per condition

**Supplemental Table 1.** Gene expression of 242 genes which were at least 2-fold upregulated in C57BL6/J lungs 3 hours after instilled ultrapure LPS exposure.

NF-κB target genes were collated from online databases (http://www.NF-kB.org, http://amp.pharm.mssm.edu/Harmonizome/gene_set/NFKB1/JASPAR+Predicted+Transcription+Factor+Targets ) and published works (http://bioinfo.lifl.fr/NF-KB , http://people.bu.edu/gilmore/nfkb/target/index.html , http://www.bu.edu/nf-kb/gene-resources/target-genes/ , References 10-12). Noncanonical NF-κB target genes were identified according to previous investigations (References 1-9).


1. T. Enzler et al., Immunity 25, 403 (2006).
2. G. Bonizzi et al., Embo J 23, 4202 (2004).
3. J. L. Chen, A. Limnander, P. B. Rothman, Blood 111, 1677 (2008
4. D. L. Drayton et al., J Immunol 173, 6161 (2004).
5. F. Guo, D. Weih, E. Meier, F. Weih, Blood 110, 2381 (2007).
6. M. Luftig et al., Proc Natl Acad Sci U S A 101, 141 (2004).
7. U. Senftleben et al., Science 293, 1495 (2001).
8. R. T. Woodland et al., Blood 111, 750 (2008).
9. J. Zhang, M. A. Warren, S. F. Shoemaker, M. M. Ip, Endocrinology 148, 268 (2007).
10. White KL, Genomics of the NF-κB signaling pathway: hypothesized role in ovarian cancerCancer Causes Control. 2011 May; 22(5): 785–801.
11. Schreiber and Jenner et al. Proc. Natl. Acad. Sci. USA 103, 2006, 5899-5904.
12. Yang Y et al, A database and functional annotation of NF-κB target genes, Int J Clin Exp Med 2016;9(5):7986-7995


| Gene | logFC WT_LPS-WT_PBS | logFC KO_LPS-KO_PBS | abslogFC (KO_LPS-KO_PBS)-(WT_LPS-WT_PBS)2 | P.Value WT_LPS-WT_PBS | P.Value KO_LPS-KO_PBS | NFkB pathway association? |
|---|---|---|---|---|---|---|
| **H2-Ea-ps** | 1.2 | 0.3 | 0.9 | 0.0164 | 0.5582 | ? |
| **Fcer2a** | 1.0 | 0.1 | 0.9 | 0.0001 | 0.7191 | canonical |
| **Cxcl2** | 7.0 | 7.8 | 0.9 | 0.0000 | 0.0000 | canonical |
| **Retnlb** | 1.2 | 2.0 | 0.8 | 0.0113 | 0.0007 | canonical |
| **Il6** | 6.5 | 7.2 | 0.7 | 0.0000 | 0.0000 | canonical |
| **Sele** | 3.3 | 4.0 | 0.7 | 0.0000 | 0.0000 | canonical |
| **Ccl12** | 3.2 | 3.9 | 0.7 | 0.0000 | 0.0000 | canonical |
| **Il23a** | 4.1 | 3.5 | 0.7 | 0.0000 | 0.0000 | canonical |
| **Camp** | 3.7 | 3.1 | 0.7 | 0.0000 | 0.0000 | canonical |
| **Elovl6** | 2.0 | 1.4 | 0.6 | 0.0000 | 0.0000 | ? |
| **Il10** | 1.2 | 0.6 | 0.6 | 0.0001 | 0.0566 | canonical |
| **Mmp12** | 1.4 | 0.8 | 0.6 | 0.0000 | 0.0001 | possible |
| **Mx1** | 5.1 | 4.5 | 0.6 | 0.0000 | 0.0000 | canonical |
| **Tnfrsf8** | 1.1 | 1.7 | 0.6 | 0.0001 | 0.0000 | canonical |
| **Cxcr2** | 4.5 | 3.9 | 0.6 | 0.0000 | 0.0000 | canonical |
| **Adamts14** | 1.1 | 0.5 | 0.6 | 0.0000 | 0.0213 | ? |

| | | | | | | |
|---|---|---|---|---|---|---|
| **Il27** | 1.0 | 1.6 | 0.6 | 0.0003 | 0.0000 | canonical |
| **Ccl4** | 5.7 | 6.2 | 0.5 | 0.0000 | 0.0000 | canonical |
| **Fcnb** | 1.6 | 1.1 | 0.5 | 0.0000 | 0.0012 | ? |
| **Elane** | 1.6 | 1.1 | 0.5 | 0.0000 | 0.0016 | canonical |
| **Cxcl11** | 5.4 | 4.9 | 0.5 | 0.0000 | 0.0000 | canonical |
| **Cxcl1** | 7.6 | 8.1 | 0.5 | 0.0000 | 0.0000 | canonical |
| **Fosl1** | 2.4 | 2.9 | 0.5 | 0.0000 | 0.0000 | ? |
| **Ptgs2** | 2.1 | 2.5 | 0.5 | 0.0000 | 0.0000 | canonical |
| **Cdkn1a** | 3.4 | 3.0 | 0.5 | 0.0000 | 0.0000 | canonical |
| **Cebpd** | 3.7 | 4.1 | 0.5 | 0.0000 | 0.0000 | possible |
| **Lif** | 1.2 | 1.6 | 0.4 | 0.0000 | 0.0000 | canonical |
| **Csf3** | 4.1 | 4.5 | 0.4 | 0.0000 | 0.0000 | canonical |
| **Nod2** | 2.8 | 2.4 | 0.4 | 0.0000 | 0.0000 | canonical |
| **Areg** | 1.4 | 1.8 | 0.4 | 0.0000 | 0.0000 | possible |
| **Ido1** | 4.2 | 3.8 | 0.4 | 0.0000 | 0.0000 | noncanonical |
| **Tlr6** | 1.5 | 1.1 | 0.4 | 0.0000 | 0.0000 | both |
| **S100a8** | 5.2 | 4.8 | 0.4 | 0.0000 | 0.0000 | predicted |
| **Cyp1b1** | 1.3 | 0.9 | 0.4 | 0.0000 | 0.0000 | ? |
| **Tnf** | 4.7 | 4.4 | 0.4 | 0.0000 | 0.0000 | canonical |
| **Crem** | 1.0 | 0.7 | 0.4 | 0.0000 | 0.0000 | canonical |
| **Ripk2** | 2.3 | 2.0 | 0.4 | 0.0000 | 0.0000 | canonical |
| **Tnfaip6** | 2.9 | 3.3 | 0.4 | 0.0000 | 0.0000 | possible |
| **Il15** | 2.6 | 2.2 | 0.4 | 0.0000 | 0.0000 | canonical |
| **Ccl24** | 1.0 | 1.4 | 0.4 | 0.0001 | 0.0000 | ? |
| **Tlr13** | 2.5 | 2.1 | 0.4 | 0.0000 | 0.0000 | ? |
| **Cd40** | 3.3 | 2.9 | 0.4 | 0.0000 | 0.0000 | canonical |
| **Mmp8** | 5.1 | 4.8 | 0.3 | 0.0000 | 0.0000 | canonical |
| **Mmp13** | 2.9 | 3.2 | 0.3 | 0.0000 | 0.0000 | possible |
| **Rgs16** | 4.2 | 4.5 | 0.3 | 0.0000 | 0.0000 | canonical |
| **Prg2** | 1.1 | 1.4 | 0.3 | 0.0011 | 0.0004 | ? |
| **Nod1** | 1.6 | 1.2 | 0.3 | 0.0000 | 0.0000 | canonical |
| **Bid** | 1.7 | 1.4 | 0.3 | 0.0000 | 0.0000 | ? |
| **S100a9** | 5.2 | 4.9 | 0.3 | 0.0000 | 0.0000 | predicted |
| **Batf** | 2.7 | 2.3 | 0.3 | 0.0000 | 0.0000 | possible |
| **Sell** | 2.2 | 1.9 | 0.3 | 0.0000 | 0.0000 | possible |
| **Lag3** | 1.5 | 1.2 | 0.3 | 0.0000 | 0.0000 | ? |
| **Irf5** | 2.0 | 1.7 | 0.3 | 0.0000 | 0.0000 | predicted |
| **Cd70** | 1.0 | 0.7 | 0.3 | 0.0003 | 0.0156 | canonical |
| **Ccl3** | 6.7 | 6.4 | 0.3 | 0.0000 | 0.0000 | canonical |
| **Casp1** | 1.5 | 1.2 | 0.3 | 0.0000 | 0.0000 | ? |
| **Il15ra** | 1.9 | 1.6 | 0.3 | 0.0000 | 0.0000 | possible |
| **Klk1** | 1.3 | 1.0 | 0.3 | 0.0000 | 0.0000 | ? |
| **Slc16a6** | 1.0 | 1.3 | 0.3 | 0.0000 | 0.0000 | ? |

| | | | | | | |
|---|---|---|---|---|---|---|
| **Mmp9** | 1.4 | 1.1 | 0.3 | 0.0000 | 0.0000 | canonical |
| **Ikbke** | 2.4 | 2.1 | 0.3 | 0.0000 | 0.0000 | canonical |
| **Tnfrsf9** | 3.7 | 3.4 | 0.3 | 0.0000 | 0.0000 | canonical |
| **P2rx1** | 1.1 | 1.4 | 0.3 | 0.0000 | 0.0000 | ? |
| **Edn1** | 1.2 | 1.0 | 0.3 | 0.0000 | 0.0000 | canonical |
| **Fcgr3** | 1.3 | 1.6 | 0.3 | 0.0000 | 0.0000 | ? |
| **Cxcl3** | 8.3 | 8.0 | 0.3 | 0.0000 | 0.0000 | canonical |
| **Irf1** | 2.4 | 2.1 | 0.3 | 0.0000 | 0.0000 | canonical |
| **Il4i1** | 5.2 | 5.5 | 0.3 | 0.0000 | 0.0000 | both |
| **Ccr7** | 1.3 | 1.0 | 0.3 | 0.0000 | 0.0000 | canonical |
| **Socs3** | 3.6 | 3.9 | 0.3 | 0.0000 | 0.0000 | ? |
| **Flnb** | 1.5 | 1.3 | 0.3 | 0.0000 | 0.0000 | ? |
| **Adora2a** | 1.7 | 1.4 | 0.3 | 0.0000 | 0.0000 | canonical |
| **Ifit1bl2** | 2.6 | 2.3 | 0.3 | 0.0000 | 0.0000 | ? |
| **Mob3c** | 2.1 | 1.8 | 0.3 | 0.0000 | 0.0000 | predicted |
| **Adamts4** | 5.3 | 5.5 | 0.3 | 0.0000 | 0.0000 | canonical |
| **Marco** | 2.2 | 1.9 | 0.3 | 0.0000 | 0.0000 | ? |
| **Clic4** | 2.3 | 2.1 | 0.3 | 0.0000 | 0.0000 | canonical |
| **Ifit1bl1** | 4.5 | 4.2 | 0.3 | 0.0000 | 0.0000 | ? |
| **Il17ra** | 1.2 | 1.0 | 0.3 | 0.0000 | 0.0000 | ? |
| **Cxcl9** | 6.3 | 6.1 | 0.3 | 0.0000 | 0.0000 | canonical |
| **Csf2** | 4.2 | 4.4 | 0.3 | 0.0000 | 0.0000 | canonical |
| **Adora3** | 1.7 | 2.0 | 0.2 | 0.0000 | 0.0000 | ? |
| **Ccl11** | 2.9 | 2.7 | 0.2 | 0.0000 | 0.0000 | canonical |
| **Cd209e** | 1.0 | 0.8 | 0.2 | 0.0005 | 0.0146 | ? |
| **Ptprc** | 1.1 | 0.8 | 0.2 | 0.0000 | 0.0000 | ? |
| **Il33** | 1.5 | 1.3 | 0.2 | 0.0000 | 0.0000 | ? |
| **Itgam** | 1.5 | 1.3 | 0.2 | 0.0000 | 0.0000 | ? |
| **Tlr9** | 1.5 | 1.3 | 0.2 | 0.0000 | 0.0000 | canonical |
| **Ptafr** | 2.9 | 2.7 | 0.2 | 0.0000 | 0.0000 | canonical |
| **Ptx3** | 6.5 | 6.7 | 0.2 | 0.0000 | 0.0000 | canonical |
| **Ccl5** | 3.1 | 2.9 | 0.2 | 0.0000 | 0.0000 | canonical |
| **Cxcl13** | 2.6 | 2.4 | 0.2 | 0.0000 | 0.0000 | predicted |
| **Gadd45b** | 2.1 | 1.9 | 0.2 | 0.0000 | 0.0000 | canonical |
| **Cd69** | 3.2 | 2.9 | 0.2 | 0.0000 | 0.0000 | canonical |
| **Nfkbiz** | 2.9 | 2.7 | 0.2 | 0.0000 | 0.0000 | canonical |
| **Sphk1** | 2.7 | 2.5 | 0.2 | 0.0000 | 0.0000 | canonical |
| **Nfil3** | 3.1 | 2.9 | 0.2 | 0.0000 | 0.0000 | possible |
| **Csf3r** | 2.6 | 2.4 | 0.2 | 0.0000 | 0.0000 | possible |
| **Ccl2** | 6.7 | 6.9 | 0.2 | 0.0000 | 0.0000 | canonical |
| **Cd80** | 2.7 | 2.5 | 0.2 | 0.0000 | 0.0000 | canonical |
| **Myd88** | 2.7 | 2.5 | 0.2 | 0.0000 | 0.0000 | canonical |
| **Tgm2** | 2.1 | 1.9 | 0.2 | 0.0000 | 0.0000 | canonical |

| | | | | | | |
|---|---|---|---|---|---|---|
| **Adam8** | 1.3 | 1.1 | 0.2 | 0.0000 | 0.0000 | ? |
| **Csf1** | 2.8 | 2.6 | 0.2 | 0.0000 | 0.0000 | canonical |
| **Maff** | 2.9 | 2.7 | 0.2 | 0.0000 | 0.0000 | ? |
| **Ly6g** | 3.4 | 3.6 | 0.2 | 0.0000 | 0.0000 | ? |
| **Birc3** | 3.3 | 3.1 | 0.2 | 0.0000 | 0.0000 | ? |
| **Klrk1** | 1.2 | 1.0 | 0.2 | 0.0000 | 0.0000 | predicted |
| **Trex1** | 3.6 | 3.4 | 0.2 | 0.0000 | 0.0000 | predicted |
| **Trafd1** | 2.7 | 2.5 | 0.2 | 0.0000 | 0.0000 | ? |
| **Gata1** | 1.0 | 0.9 | 0.2 | 0.0000 | 0.0003 | ? |
| **Tap2** | 1.6 | 1.4 | 0.2 | 0.0000 | 0.0000 | predicted |
| **Birc2** | 2.2 | 2.0 | 0.2 | 0.0000 | 0.0000 | canonical |
| **H2-M3** | 1.1 | 0.9 | 0.2 | 0.0000 | 0.0000 | ? |
| **Adamts9** | 2.5 | 2.3 | 0.2 | 0.0000 | 0.0000 | ? |
| **Hif1a** | 2.1 | 1.9 | 0.2 | 0.0000 | 0.0000 | canonical |
| **Casp7** | 1.0 | 0.8 | 0.2 | 0.0000 | 0.0000 | predicted |
| **Rin2** | 1.0 | 0.8 | 0.2 | 0.0000 | 0.0000 | predicted |
| **Tnfaip3** | 4.6 | 4.4 | 0.2 | 0.0000 | 0.0000 | canonical |
| **Malt1** | 2.7 | 2.5 | 0.2 | 0.0000 | 0.0000 | predicted |
| **Il4ra** | 2.2 | 2.0 | 0.2 | 0.0000 | 0.0000 | ? |
| **Nfkbie** | 3.5 | 3.4 | 0.2 | 0.0000 | 0.0000 | canonical |
| **Pdcd1** | 2.0 | 1.8 | 0.2 | 0.0000 | 0.0000 | canonical |
| **Batf3** | 1.3 | 1.4 | 0.2 | 0.0000 | 0.0000 | predicted |
| **Cyr61** | 1.2 | 1.0 | 0.2 | 0.0000 | 0.0000 | ? |
| **Ccrl2** | 3.1 | 2.9 | 0.2 | 0.0000 | 0.0000 | predicted |
| **Ccl7** | 5.4 | 5.6 | 0.2 | 0.0000 | 0.0000 | predicted |
| **Mx2** | 5.5 | 5.6 | 0.2 | 0.0000 | 0.0000 | ? |
| **Fcgr1** | 3.1 | 3.0 | 0.2 | 0.0000 | 0.0000 | ? |
| **Amica1** | 2.5 | 2.3 | 0.2 | 0.0000 | 0.0000 | ? |
| **Nampt** | 2.4 | 2.2 | 0.2 | 0.0000 | 0.0000 | possible |
| **Stat1** | 2.0 | 1.8 | 0.2 | 0.0000 | 0.0000 | predicted |
| **Fscn1** | 1.2 | 1.0 | 0.2 | 0.0000 | 0.0000 | ? |
| **Cd274** | 4.5 | 4.3 | 0.2 | 0.0000 | 0.0000 | canonical |
| **Fpr1** | 2.0 | 2.2 | 0.2 | 0.0000 | 0.0000 | ? |
| **Cd47** | 1.1 | 1.0 | 0.2 | 0.0000 | 0.0000 | ? |
| **Plaur** | 2.1 | 2.2 | 0.2 | 0.0000 | 0.0000 | ? |
| **Ccl17** | 4.5 | 4.6 | 0.2 | 0.0000 | 0.0000 | canonical |
| **Serpine1** | 4.7 | 4.5 | 0.2 | 0.0000 | 0.0000 | ? |
| **Stat3** | 1.4 | 1.3 | 0.2 | 0.0000 | 0.0000 | possible |
| **Selp** | 4.6 | 4.7 | 0.1 | 0.0000 | 0.0000 | canonical |
| **Vcam1** | 2.8 | 3.0 | 0.1 | 0.0000 | 0.0000 | canonical |
| **Tnfrsf1b** | 2.0 | 1.8 | 0.1 | 0.0000 | 0.0000 | canonical |
| **Psmb9** | 1.3 | 1.1 | 0.1 | 0.0000 | 0.0000 | canonical |
| **Tnc** | 1.0 | 1.1 | 0.1 | 0.0000 | 0.0000 | canonical |

| | | | | | | |
|---|---|---|---|---|---|---|
| **Nfkbia** | 3.3 | 3.2 | 0.1 | 0.0000 | 0.0000 | canonical |
| **Tlr3** | 2.1 | 1.9 | 0.1 | 0.0000 | 0.0000 | ? |
| **Tnfsf10** | 1.2 | 1.1 | 0.1 | 0.0000 | 0.0000 | canonical |
| **Il12b** | 2.3 | 2.5 | 0.1 | 0.0000 | 0.0000 | canonical |
| **Tlr7** | 1.3 | 1.2 | 0.1 | 0.0000 | 0.0000 | ? |
| **Fcgr2b** | 1.9 | 1.7 | 0.1 | 0.0000 | 0.0000 | possible |
| **Cd14** | 5.4 | 5.3 | 0.1 | 0.0000 | 0.0000 | possible |
| **Hdc** | 2.6 | 2.5 | 0.1 | 0.0000 | 0.0000 | ? |
| **Cebpb** | 1.7 | 1.6 | 0.1 | 0.0000 | 0.0000 | predicted |
| **Gem** | 2.6 | 2.7 | 0.1 | 0.0000 | 0.0000 | ? |
| **Fut4** | 1.3 | 1.1 | 0.1 | 0.0000 | 0.0000 | ? |
| **Il1a** | 4.7 | 4.6 | 0.1 | 0.0000 | 0.0000 | canonical |
| **Pglyrp1** | 1.9 | 2.0 | 0.1 | 0.0000 | 0.0000 | canonical |
| **Fcgr4** | 2.1 | 2.0 | 0.1 | 0.0000 | 0.0000 | possible |
| **Tmem173** | 1.9 | 2.0 | 0.1 | 0.0000 | 0.0000 | predicted |
| **Emp1** | 1.3 | 1.2 | 0.1 | 0.0000 | 0.0000 | ? |
| **Bcl10** | 1.3 | 1.2 | 0.1 | 0.0000 | 0.0000 | possible |
| **Il10ra** | 1.6 | 1.5 | 0.1 | 0.0000 | 0.0000 | ? |
| **Ccl9** | 3.1 | 3.0 | 0.1 | 0.0000 | 0.0000 | ? |
| **Hbegf** | 1.7 | 1.8 | 0.1 | 0.0000 | 0.0000 | ? |
| **Map2k1** | 1.1 | 1.0 | 0.1 | 0.0000 | 0.0000 | ? |
| **Saa1** | 5.1 | 5.0 | 0.1 | 0.0000 | 0.0000 | canonical |
| **Ccl22** | 4.4 | 4.5 | 0.1 | 0.0000 | 0.0000 | predicted |
| **Tapbp** | 1.4 | 1.3 | 0.1 | 0.0000 | 0.0000 | canonical |
| **Ccr5** | 1.0 | 1.1 | 0.1 | 0.0000 | 0.0000 | canonical |
| **Stat5a** | 1.4 | 1.3 | 0.1 | 0.0000 | 0.0000 | canonical |
| **Irf2** | 1.3 | 1.2 | 0.1 | 0.0000 | 0.0000 | canonical |
| **H2-K1** | 1.2 | 1.1 | 0.1 | 0.0000 | 0.0000 | ? |
| **Cd86** | 2.3 | 2.4 | 0.1 | 0.0000 | 0.0000 | canonical |
| **Tnfrsf12a** | 2.2 | 2.3 | 0.1 | 0.0000 | 0.0000 | canonical |
| **S1pr1** | 1.6 | 1.5 | 0.1 | 0.0000 | 0.0000 | ? |
| **Ell2** | 1.0 | 0.9 | 0.1 | 0.0000 | 0.0000 | predicted |
| **Ccr1** | 3.0 | 3.1 | 0.1 | 0.0000 | 0.0000 | ? |
| **Psmb8** | 1.8 | 1.7 | 0.1 | 0.0000 | 0.0000 | ? |
| **Prok2** | 1.8 | 1.9 | 0.1 | 0.0000 | 0.0000 | ? |
| **Il25** | 1.0 | 1.0 | 0.1 | 0.0037 | 0.0039 | predicted |
| **Rhoj** | 1.4 | 1.3 | 0.1 | 0.0000 | 0.0000 | ? |
| **Cxcl5** | 6.5 | 6.6 | 0.1 | 0.0000 | 0.0000 | canonical |
| **Itga5** | 1.0 | 0.9 | 0.1 | 0.0000 | 0.0000 | predicted |
| **Tap1** | 3.1 | 3.0 | 0.1 | 0.0000 | 0.0000 | canonical |
| **Rgs1** | 2.7 | 2.8 | 0.1 | 0.0000 | 0.0000 | ? |
| **Irf7** | 4.5 | 4.5 | 0.1 | 0.0000 | 0.0000 | canonical |
| **Cd44** | 1.5 | 1.5 | 0.1 | 0.0000 | 0.0000 | canonical |

| | | | | | | |
|---|---|---|---|---|---|---|
| **H2-Q1** | 1.4 | 1.3 | 0.1 | 0.0000 | 0.0000 | ? |
| **Enc1** | 1.4 | 1.3 | 0.1 | 0.0000 | 0.0000 | predicted |
| **Osm** | 3.6 | 3.5 | 0.1 | 0.0000 | 0.0000 | ? |
| **Cx3cl1** | 1.1 | 1.0 | 0.1 | 0.0000 | 0.0000 | ? |
| **Nfkb1** | 1.4 | 1.3 | 0.1 | 0.0000 | 0.0000 | canonical |
| **C5ar1** | 1.9 | 1.8 | 0.1 | 0.0000 | 0.0000 | ? |
| **Ier3** | 1.2 | 1.2 | 0.1 | 0.0000 | 0.0000 | canonical |
| **H2-D1** | 1.1 | 1.0 | 0.1 | 0.0000 | 0.0000 | ? |
| **Anxa1** | 2.4 | 2.3 | 0.1 | 0.0000 | 0.0000 | ? |
| **S100a11** | 1.0 | 0.9 | 0.1 | 0.0000 | 0.0000 | ? |
| **Cybb** | 2.0 | 1.9 | 0.1 | 0.0000 | 0.0000 | possible |
| **Tlr2** | 3.4 | 3.4 | 0.1 | 0.0000 | 0.0000 | ? |
| **Nlrp3** | 3.7 | 3.7 | 0.1 | 0.0000 | 0.0000 | canonical |
| **Gch1** | 2.5 | 2.4 | 0.1 | 0.0000 | 0.0000 | ? |
| **Aoah** | 2.3 | 2.3 | 0.1 | 0.0000 | 0.0000 | ? |
| **Peli1** | 1.0 | 1.0 | 0.1 | 0.0000 | 0.0000 | ? |
| **Cxcl10** | 10.3 | 10.3 | 0.1 | 0.0000 | 0.0000 | canonical |
| **Syk** | 1.2 | 1.2 | 0.1 | 0.0000 | 0.0000 | ? |
| **Clec5a** | 2.6 | 2.6 | 0.1 | 0.0000 | 0.0000 | ? |
| **Ceacam1** | 1.5 | 1.4 | 0.1 | 0.0000 | 0.0000 | ? |
| **Timd4** | 1.7 | 1.7 | 0.1 | 0.0000 | 0.0000 | ? |
| **Ccl20** | 8.3 | 8.2 | 0.1 | 0.0000 | 0.0000 | canonical |
| **Usp18** | 4.5 | 4.4 | 0.0 | 0.0000 | 0.0000 | canonical |
| **Il1b** | 6.1 | 6.2 | 0.0 | 0.0000 | 0.0000 | canonical |
| **Tnfsf9** | 1.4 | 1.4 | 0.0 | 0.0000 | 0.0000 | canonical |
| **Icosl** | 1.3 | 1.3 | 0.0 | 0.0000 | 0.0000 | ? |
| **Ly6c1** | 1.5 | 1.5 | 0.0 | 0.0000 | 0.0000 | ? |
| **Isg15** | 5.6 | 5.6 | 0.0 | 0.0000 | 0.0000 | ? |
| **Cxcl16** | 1.9 | 1.9 | 0.0 | 0.0000 | 0.0000 | predicted |
| **Rnd3** | 1.5 | 1.5 | 0.0 | 0.0000 | 0.0000 | ? |
| **H2-Q10** | 1.8 | 1.8 | 0.0 | 0.0000 | 0.0000 | ? |
| **Trem1** | 3.0 | 3.1 | 0.0 | 0.0000 | 0.0000 | canonical |
| **Smad1** | 1.2 | 1.2 | 0.0 | 0.0000 | 0.0000 | ? |
| **Il1r1** | 1.1 | 1.1 | 0.0 | 0.0000 | 0.0000 | ? |
| **Icam1** | 2.7 | 2.7 | 0.0 | 0.0000 | 0.0000 | canonical |
| **Cd83** | 3.2 | 3.2 | 0.0 | 0.0000 | 0.0000 | canonical |
| **Tnfrsf11a** | 1.9 | 1.9 | 0.0 | 0.0000 | 0.0000 | ? |
| **Fpr2** | 3.4 | 3.4 | 0.0 | 0.0000 | 0.0000 | ? |
| **Daxx** | 3.1 | 3.1 | 0.0 | 0.0000 | 0.0000 | ? |
| **Socs1** | 3.1 | 3.1 | 0.0 | 0.0000 | 0.0000 | ? |
| **Atf3** | 3.5 | 3.5 | 0.0 | 0.0000 | 0.0000 | predicted |
| **Gpr65** | 1.6 | 1.6 | 0.0 | 0.0000 | 0.0000 | ? |
| **S100a10** | 1.2 | 1.2 | 0.0 | 0.0000 | 0.0000 | canonical |

| | | | | | | |
|---|---|---|---|---|---|---|
| **Psme2** | 1.7 | 1.7 | 0.0 | 0.0000 | 0.0000 | canonical |
| **Dusp1** | 1.7 | 1.7 | 0.0 | 0.0000 | 0.0000 | canonical |
| **Marcksl1** | 4.5 | 4.5 | 0.0 | 0.0000 | 0.0000 | ? |
| **Acod1** | 6.1 | 6.1 | 0.0 | 0.0000 | 0.0000 | ? |
| **Rab20** | 2.9 | 2.9 | 0.0 | 0.0000 | 0.0000 | ? |
| **Ccl19** | 4.6 | 4.5 | 0.0 | 0.0000 | 0.0000 | canonical |
| **Fas** | 2.7 | 2.7 | 0.0 | 0.0000 | 0.0000 | canonical |
| **Dusp2** | 3.3 | 3.3 | 0.0 | 0.0000 | 0.0000 | predicted |
| **Sema4a** | 1.1 | 1.1 | 0.0 | 0.0000 | 0.0000 | predicted |
| **Rhoc** | 2.2 | 2.2 | 0.0 | 0.0000 | 0.0000 | predicted |
| **Ctla4** | 1.3 | 1.3 | 0.0 | 0.0000 | 0.0001 | ? |

| **logFCWT-KO** | **% NFkB responsive genes** | **All genes** | **not predicted NFkB responsive** | **predicted NFkB responsive** | | **Notes** |
|---|---|---|---|---|---|---|
| 0.7-0.9 | **88.9** | 9 | 1 | 8 | | 62%-86% difference between KO and WT |
| 0.5-0.6 | **76.5** | 17 | 4 | 13 | | 41%-52% difference between KO and WT |
| 0.4 | **81.3** | 16 | 3 | 13 | | 32% difference between KO and WT |
| 0.3 | **64.1** | 39 | 14 | 25 | | 25% difference between KO and WT |
| 0.2 | **61.0** | 59 | 23 | 36 | | 15% difference between KO and WT |
| 0.1 | **54.3** | 70 | 32 | 38 | | 7% difference between KO and WT |
| 0 | **50.0** | 32 | 16 | 16 | | no difference |

**Supplemental Table 2.** Gene expression of all genes analyzed in C57BL6/J lungs 3 hours after instilled PBS exposure.

| Gene | GeneClass | adj.P.Val KO_PBS-WT_PBS |
|---|---|---|
| Il4 | | 0.83 |
| Cx3cr1 | | 1.00 |
| Ear3 | | 1.00 |
| Ccl28 | | 1.00 |
| Nfatc2 | | 1.00 |
| Ccr7 | | 1.00 |
| Ccl22 | | 1.00 |
| Tnfaip3 | | 1.00 |
| Ptprb | | 1.00 |
| Ccl21a | | 1.00 |
| Trim9 | | 1.00 |
| Olr1 | | 1.00 |
| Ets1 | | 1.00 |
| Serpine1 | | 1.00 |
| Stat5a | | 1.00 |
| Flrt2 | | 1.00 |
| Hip1r | | 1.00 |
| Ndufa7 | | 1.00 |
| Ctla4 | | 1.00 |
| Mob3c | | 1.00 |
| Fgf2 | | 1.00 |
| Mok | | 0.84 |
| Ccl26 | | 1.00 |
| Cd68 | | 1.00 |
| Hras | | 1.00 |
| Vav2 | | 1.00 |
| C3ar1 | | 1.00 |
| Btk | | 1.00 |
| Ccl4 | | 1.00 |
| Crem | | 1.00 |
| Fcer1a | | 1.00 |
| Cd40 | | 1.00 |
| Twistnb | | 1.00 |
| H2-Eb1 | | 1.00 |
| Cd86 | | 1.00 |
| Tlr9 | | 1.00 |
| Il1rl1 | | 1.00 |
| Psme2 | | 1.00 |
| Fcnb | | 1.00 |
| Nfil3 | | 1.00 |
| Clec5a | | 1.00 |

| | | |
|---|---|---|
| **Cma1** | | 0.83 |
| **Ski** | | 1.00 |
| **Nox1** | | 0.83 |
| **Hc** | | 1.00 |
| **Igf2** | | 1.00 |
| **Pdzk1ip1** | | 1.00 |
| **Nr4a2** | | 1.00 |
| **Crip1** | | 1.00 |
| **Hdc** | | 1.00 |
| **Bmp8a** | | 1.00 |
| **Ccl24** | | 1.00 |
| **Ccl20** | | 1.00 |
| **Cd274** | | 1.00 |
| **Des** | | 1.00 |
| **Ikzf1** | | 1.00 |
| **Cd80** | | 1.00 |
| **Ccr3** | | 1.00 |
| **Ccl3** | | 1.00 |
| **Lpl** | | 0.83 |
| **Irf4** | | 1.00 |
| **Klf10** | | 1.00 |
| **Hif1a** | | 1.00 |
| **Epx** | | 1.00 |
| **Il12b** | | 0.83 |
| **Fem1c** | | 1.00 |
| **Fcgr1** | | 1.00 |
| **Klk1** | | 1.00 |
| **Il4ra** | | 1.00 |
| **H2-Q10** | | 1.00 |
| **Raf1** | | 1.00 |
| **Flt1** | | 1.00 |
| **Adamts1** | | 1.00 |
| **Tmem173** | | 1.00 |
| **Kif20a** | | 1.00 |
| **Socs3** | | 1.00 |
| **Glg1** | | 1.00 |
| **Serpine3** | | 1.00 |
| **Sirpa** | | 1.00 |
| **Acox1** | | 1.00 |
| **Adamts17** | | 1.00 |
| **Havcr1** | | 1.00 |
| **Gadd45b** | | 1.00 |
| **Cxcr1** | | 1.00 |
| **H2-Q2** | | 1.00 |
| **Cxcl2** | | 1.00 |
| **Col11a1** | | 1.00 |
| **Il3ra** | | 0.83 |
| **Emp1** | | 1.00 |

| | | |
|---|---|---|
| Tgfb1 | | 1.00 |
| Vwa5a | | 1.00 |
| P2rx1 | | 1.00 |
| Fasn | | 1.00 |
| S100a4 | | 1.00 |
| Tnfrsf14 | | 1.00 |
| H2-DMa | | 1.00 |
| Adamts4 | | 1.00 |
| Mapk14 | | 1.00 |
| Cxcl10 | | 1.00 |
| Pf4 | | 1.00 |
| Sgpp1 | | 1.00 |
| Lat2 | | 1.00 |
| Fbxl7 | | 1.00 |
| Klrk1 | | 1.00 |
| Gch1 | | 1.00 |
| Ctnnb1 | | 1.00 |
| Amica1 | | 1.00 |
| Prg2 | | 1.00 |
| Ros1 | | 1.00 |
| Cd247 | | 1.00 |
| Mpo | | 1.00 |
| Gzma | | 1.00 |
| Tep1 | | 1.00 |
| Hnf1b | | 1.00 |
| Mmp10 | | 1.00 |
| Prg3 | | 1.00 |
| Fbp1 | | 1.00 |
| Runx2 | | 1.00 |
| Cybb | | 1.00 |
| Nlrp3 | | 1.00 |
| Cytip | | 1.00 |
| Ctsg | | 1.00 |
| Nmb | | 1.00 |
| Cd36 | | 1.00 |
| Cxcl13 | | 1.00 |
| Tapbp | | 1.00 |
| Ier3 | | 1.00 |
| Ifng | | 1.00 |
| Stat6 | | 1.00 |
| Tap1 | | 1.00 |
| Myh4 | | 1.00 |
| Man2b1 | | 1.00 |
| Trem2 | | 1.00 |
| Hist2h2aa1 | | 1.00 |
| Il2 | | 1.00 |
| Pfdn6 | | 1.00 |
| Cdh4 | | 1.00 |

| | | |
|---|---|---|
| **Csf1r** | | 1.00 |
| **Gnai3** | | 1.00 |
| **Lat** | | 1.00 |
| **Jun** | | 1.00 |
| **Il1b** | | 1.00 |
| **Tgm2** | | 1.00 |
| **Cd163** | | 0.83 |
| **Cxcr4** | | 1.00 |
| **Vtcn1** | | 1.00 |
| **S100a11** | | 1.00 |
| **Ccr9** | | 1.00 |
| **Il15** | | 1.00 |
| **Ptger2** | | 1.00 |
| **C5ar1** | | 1.00 |
| **Yes1** | | 1.00 |
| **Hdac6** | | 1.00 |
| **Alox5** | | 1.00 |
| **Oscar** | | 1.00 |
| **Cd180** | | 1.00 |
| **Ifit1bl2** | | 1.00 |
| **Rhog** | | 1.00 |
| **Arf6** | | 1.00 |
| **Itgax** | | 1.00 |
| **Ell2** | | 1.00 |
| **Dusp6** | | 1.00 |
| **Vegfc** | | 1.00 |
| **Tlr1** | | 1.00 |
| **Daxx** | | 1.00 |
| **Cxcl3** | | 1.00 |
| **Il18r1** | | 1.00 |
| **Retnlb** | | 1.00 |
| **Selplg** | | 1.00 |
| **Igf1** | | 1.00 |
| **Cd69** | | 1.00 |
| **Cd1d1** | | 1.00 |
| **H2-Ob** | | 1.00 |
| **Loxl2** | | 1.00 |
| **Lrp5** | | 1.00 |
| **Id1** | | 1.00 |
| **Il33** | | 1.00 |
| **Il10ra** | | 1.00 |
| **2810417H13Rik** | | 1.00 |
| **Ccr1** | | 1.00 |
| **Mx1** | | 1.00 |
| **Acly** | | 1.00 |
| **H2-Ea-ps** | | 1.00 |
| **Clec1b** | | 1.00 |
| **Saa1** | | 1.00 |

| | | |
|---|---|---|
| **Dpp4** | | 1.00 |
| **Ripk2** | | 1.00 |
| **H2-Aa** | | 1.00 |
| **Tnfrsf9** | | 1.00 |
| **Lif** | | 1.00 |
| **Gk5** | | 1.00 |
| **Fyn** | | 1.00 |
| **Syk** | | 1.00 |
| **Il23a** | | 0.83 |
| **Col12a1** | | 1.00 |
| **Gpr183** | | 1.00 |
| **Tm7sf3** | | 1.00 |
| **B4galt4** | | 1.00 |
| **H2-DMb1** | | 1.00 |
| **Ephb6** | | 1.00 |
| **Marcksl1** | | 1.00 |
| **Flt3** | | 1.00 |
| **Fas** | | 1.00 |
| **Csf1** | | 1.00 |
| **Il25** | | 1.00 |
| **Birc2** | | 1.00 |
| **Tnfrsf12a** | | 1.00 |
| **Prkci** | | 1.00 |
| **Irf7** | | 1.00 |
| **Arhgef6** | | 1.00 |
| **Nfatc1** | | 1.00 |
| **Vamp2** | | 1.00 |
| **H2-Q1** | | 1.00 |
| **Cd1d2** | | 1.00 |
| **Nos2** | | 1.00 |
| **Stat5b** | | 1.00 |
| **Birc5** | | 1.00 |
| **Adam8** | | 1.00 |
| **Osm** | | 1.00 |
| **Alcam** | | 1.00 |
| **Serpinb6a** | | 1.00 |
| **Apoe** | | 0.86 |
| **Ccl6** | | 1.00 |
| **Fscn1** | | 1.00 |
| **S100a8** | | 1.00 |
| **Maff** | | 1.00 |
| **Gata1** | | 1.00 |
| **Krba1** | | 1.00 |
| **Fgf18** | | 1.00 |
| **Plau** | | 1.00 |
| **Adgre1** | | 1.00 |
| **Ccl17** | | 1.00 |
| **Cdc20** | | 1.00 |

| | | |
|---|---|---|
| Casp1 | | 1.00 |
| Cxcl14 | | 1.00 |
| Adamts2 | | 0.83 |
| Cd74 | | 0.95 |
| Jag1 | | 1.00 |
| C3 | | 1.00 |
| Ttk | | 1.00 |
| Traf2 | | 1.00 |
| Socs1 | | 1.00 |
| Calr | | 1.00 |
| Ctsd | | 1.00 |
| Tnfaip8 | | 1.00 |
| H2-T23 | | 1.00 |
| Prdx3 | | 1.00 |
| Rab3d | | 1.00 |
| Siglec1 | | 1.00 |
| Aoah | | 1.00 |
| Hoxd4 | | 1.00 |
| Irf3 | | 1.00 |
| Sele | | 1.00 |
| Ccr4 | | 1.00 |
| Cdh5 | | 1.00 |
| Map2k3 | | 1.00 |
| Ctsa | | 1.00 |
| Irf2 | | 1.00 |
| Gsn | | 0.83 |
| Clec9a | | 1.00 |
| Hes1 | | 1.00 |
| Il12a | | 1.00 |
| Il5 | | 1.00 |
| Tspan7 | | 1.00 |
| Cebpg | | 1.00 |
| Cdh1 | | 1.00 |
| Pbx3 | | 1.00 |
| Smad1 | | 1.00 |
| Tspan8 | | 1.00 |
| Cd70 | | 1.00 |
| Ptprc | | 1.00 |
| Tuba4a | | 1.00 |
| Sesn1 | | 1.00 |
| Lamb3 | | 1.00 |
| Il1r2 | | 1.00 |
| Vsir | | 1.00 |
| Il1rap | | 1.00 |
| Ccr6 | | 1.00 |
| Fpr-rs3 | | 0.83 |
| Ccl5 | | 1.00 |
| Nod1 | | 1.00 |

| | | |
|---|---|---|
| **Cdh2** | | 1.00 |
| **Il6ra** | | 1.00 |
| **Irf1** | | 1.00 |
| **Cx3cl1** | | 1.00 |
| **Mx2** | | 1.00 |
| **H2-DMb2** | | 1.00 |
| **Fzd9** | | 1.00 |
| **Cd14** | | 1.00 |
| **Ccl25** | | 1.00 |
| **Zfp148** | | 1.00 |
| **Tlr8** | | 1.00 |
| **Fpr-rs6** | | 1.00 |
| **Pdgfa** | | 1.00 |
| **Cebpa** | | 1.00 |
| **Gpr65** | | 1.00 |
| **Fpr3** | | 0.83 |
| **Csf3** | | 1.00 |
| **Timd4** | | 1.00 |
| **Id2** | | 1.00 |
| **Txndc16** | | 1.00 |
| **Camp** | | 1.00 |
| **Tlr11** | | 1.00 |
| **Pim2** | | 1.00 |
| **Cd28** | | 1.00 |
| **Znrf2** | | 1.00 |
| **Col15a1** | | 0.83 |
| **Adora3** | | 1.00 |
| **Ccl9** | | 1.00 |
| **Cxcl16** | | 1.00 |
| **Cbr1** | | 1.00 |
| **Ptprk** | | 1.00 |
| **Fcgr4** | | 1.00 |
| **Il4i1** | | 0.83 |
| **Myod1** | | 1.00 |
| **Rgs1** | | 1.00 |
| **Mapk13** | | 1.00 |
| **Erg** | | 1.00 |
| **Bid** | | 1.00 |
| **Rasgrf2** | | 1.00 |
| **Ly6g** | | 0.95 |
| **Ccr8** | | 1.00 |
| **H2-D1** | | 1.00 |
| **Ceacam1** | | 1.00 |
| **Bmp6** | | 1.00 |
| **Col4a1** | | 1.00 |
| **Myc** | | 1.00 |
| **Cd27** | | 1.00 |
| **Cxcl12** | | 1.00 |

| | | |
|---|---|---|
| **Phlda2** | | 1.00 |
| **Faf1** | | 1.00 |
| **Sqstm1** | | 1.00 |
| **Vrk2** | | 1.00 |
| **Cd34** | | 0.93 |
| **Furin** | | 1.00 |
| **Nfkb1** | | 1.00 |
| **Smad2** | | 1.00 |
| **Cxcl1** | | 1.00 |
| **Casp7** | | 1.00 |
| **Icos** | | 1.00 |
| **Gata3** | | 1.00 |
| **Ccl8** | | 0.37 |
| **Fut4** | | 1.00 |
| **Il27ra** | | 1.00 |
| **Map3k4** | | 1.00 |
| **Fpr-rs5** | | 1.00 |
| **Nampt** | | 1.00 |
| **H2-M3** | | 1.00 |
| **Ceacam2** | | 1.00 |
| **Usp18** | | 1.00 |
| **Tnf** | | 1.00 |
| **Tfcp2** | | 1.00 |
| **Lrp6** | | 1.00 |
| **Serpinb9** | | 1.00 |
| **Msh2** | | 1.00 |
| **Sfrp4** | | 0.83 |
| **Cysltr1** | | 1.00 |
| **Crabp2** | | 1.00 |
| **S1pr1** | | 1.00 |
| **Tnfrsf1a** | | 1.00 |
| **Atf3** | | 1.00 |
| **Top2a** | | 1.00 |
| **Tlr7** | | 1.00 |
| **Cd151** | | 1.00 |
| **Col14a1** | | 0.07 |
| **Lamb2** | | 1.00 |
| **Col17a1** | | 1.00 |
| **C1qc** | | 0.37 |
| **Rgl1** | | 1.00 |
| **Dnajb6** | | 1.00 |
| **Eif4ebp2** | | 1.00 |
| **Il1a** | | 1.00 |
| **Ltb** | | 1.00 |
| **Icam1** | | 1.00 |
| **Cxcl9** | | 1.00 |
| **Mrc2** | | 1.00 |
| **Mmp1a** | | 1.00 |

| | | |
|---|---|---|
| **Angpt1** | | 1.00 |
| **Ptgs2** | | 0.83 |
| **Alox5ap** | | 1.00 |
| **Pdgfrb** | | 1.00 |
| **Cyr61** | | 1.00 |
| **Txn1** | | 1.00 |
| **Adamts12** | | 1.00 |
| **Rasal1** | | 1.00 |
| **Itgal** | | 1.00 |
| **Cxcl11** | | 1.00 |
| **Ccrl2** | | 1.00 |
| **Egf** | | 1.00 |
| **Ptgdr** | | 0.83 |
| **Nrip3** | | 1.00 |
| **Pdcd1lg2** | | 1.00 |
| **Cav1** | | 1.00 |
| **Acad10** | | 1.00 |
| **Cnksr1** | | 1.00 |
| **Tgfbr3** | | 1.00 |
| **Stat1** | | 1.00 |
| **Hist1h1c** | | 1.00 |
| **Chil3** | | 1.00 |
| **Pdgfb** | | 1.00 |
| **Egr3** | | 1.00 |
| **Enpep** | | 1.00 |
| **Slc16a6** | | 1.00 |
| **Il13** | | 1.00 |
| **Pros1** | | 1.00 |
| **Adcyap1r1** | | 1.00 |
| **Ccl11** | | 1.00 |
| **Cldn1** | | 1.00 |
| **Lgals3** | | 1.00 |
| **Rapgef4** | | 1.00 |
| **Col1a2** | | 0.83 |
| **Enpp2** | | 0.99 |
| **Btla** | | 1.00 |
| **Spry2** | | 1.00 |
| **Map3k14** | | 1.00 |
| **Serpinb2** | | 1.00 |
| **Zmpste24** | | 1.00 |
| **Sell** | | 1.00 |
| **Cd47** | | 1.00 |
| **Nr4a1** | | 1.00 |
| **Enc1** | | 1.00 |
| **Mapk11** | | 1.00 |
| **Fosl1** | | 1.00 |
| **Hbegf** | | 1.00 |
| **Fpr2** | | 1.00 |

| | | |
|---|---|---|
| **Cdh13** | | 0.37 |
| **Prkca** | | 1.00 |
| **Ptpn14** | | 1.00 |
| **Rhoc** | | 1.00 |
| **Tlr5** | | 0.00 |
| **Adamts3** | | 1.00 |
| **Cd38** | | 1.00 |
| **Jak3** | | 1.00 |
| **Rnase2a** | | 0.83 |
| **Cd244** | | 1.00 |
| **Ctss** | | 1.00 |
| **Lst1** | | 1.00 |
| **Itgam** | | 1.00 |
| **Grn** | | 0.89 |
| **Il3** | | 1.00 |
| **Malt1** | | 1.00 |
| **Angpt2** | | 1.00 |
| **Ly6c1** | | 1.00 |
| **Elovl6** | | 0.84 |
| **Tnfsf9** | | 1.00 |
| **Hivep1** | | 1.00 |
| **Areg** | | 0.83 |
| **Fpr-rs4** | | 1.00 |
| **Cd99** | | 1.00 |
| **Fosb** | | 1.00 |
| **Cd276** | | 1.00 |
| **Skil** | | 1.00 |
| **Dusp1** | | 0.83 |
| **Tlr2** | | 1.00 |
| **Cttnbp2** | | 1.00 |
| **Tlr6** | | 1.00 |
| **Thbd** | | 1.00 |
| **F11r** | | 1.00 |
| **Btg2** | | 1.00 |
| **Clic4** | | 1.00 |
| **Il18** | | 1.00 |
| **Golim4** | | 1.00 |
| **Bcl2** | | 1.00 |
| **Chil4** | | 1.00 |
| **Kcnq1ot1** | | 1.00 |
| **Rad51** | | 1.00 |
| **Tnc** | | 1.00 |
| **Tlr3** | | 1.00 |
| **Ndc80** | | 0.37 |
| **Ngf** | | 1.00 |
| **Aif1** | | 0.37 |
| **Pparg** | | 1.00 |
| **Egr2** | | 1.00 |

| | | |
|---|---|---|
| **Fzd6** | | 1.00 |
| **Gata2** | | 1.00 |
| **Mcm5** | | 1.00 |
| **Tnfsf4** | | 1.00 |
| **Cxcr5** | | 1.00 |
| **Mmp9** | | 1.00 |
| **Ccl2** | | 1.00 |
| **Acot3** | | 1.00 |
| **Nr1h3** | | 1.00 |
| **Il21r** | | 1.00 |
| **Ms4a1** | | 1.00 |
| **Rnd3** | | 1.00 |
| **Traf1** | | 1.00 |
| **Prdx1** | | 1.00 |
| **Ifnb1** | | 1.00 |
| **Rnase2b** | | 1.00 |
| **Cstb** | | 1.00 |
| **Dhrs3** | | 1.00 |
| **Nod2** | | 1.00 |
| **Yap1** | | 1.00 |
| **Ncor1** | | 1.00 |
| **H2-K1** | | 1.00 |
| **Irf5** | | 1.00 |
| **Bcl6** | | 1.00 |
| **Met** | | 1.00 |
| **Hpgds** | | 1.00 |
| **Nr2f6** | | 1.00 |
| **Nfkbiz** | | 1.00 |
| **Selp** | | 1.00 |
| **Tek** | | 1.00 |
| **Mafb** | | 1.00 |
| **C1qa** | | 0.37 |
| **Vegfa** | | 1.00 |
| **Anxa4** | | 1.00 |
| **Mmp13** | | 1.00 |
| **Lamp3** | | 1.00 |
| **Stab1** | | 1.00 |
| **Rbp1** | | 1.00 |
| **Hdac5** | | 1.00 |
| **Cxcr6** | | 1.00 |
| **Kalrn** | | 1.00 |
| **Arg1** | | 1.00 |
| **Il9** | | 1.00 |
| **Laptm5** | | 1.00 |
| **Smad7** | | 1.00 |
| **Ptgdr2** | | 1.00 |
| **Il15ra** | | 1.00 |
| **Tgfa** | | 1.00 |

| | | |
|---|---|---|
| **Tnfrsf4** | | 1.00 |
| **Cnn2** | | 1.00 |
| **Ptgds** | | 0.83 |
| **Csf2** | | 1.00 |
| **Ifnar2** | | 0.83 |
| **Flnb** | | 1.00 |
| **Ercc1** | | 1.00 |
| **Ccl27a** | | 1.00 |
| **Mmp19** | | 1.00 |
| **Sema4b** | | 1.00 |
| **Vav1** | | 1.00 |
| **Ncf2** | | 1.00 |
| **Tnfrsf11a** | | 1.00 |
| **Lta4h** | | 1.00 |
| **Itgb7** | | 1.00 |
| **Hgf** | | 1.00 |
| **Cebpd** | | 1.00 |
| **Itgb2** | | 1.00 |
| **Pik3cg** | | 1.00 |
| **Cyp3a13** | | 1.00 |
| **Batf3** | | 1.00 |
| **S100a9** | | 1.00 |
| **Psmb8** | | 1.00 |
| **Acod1** | | 1.00 |
| **Ifna1** | | 1.00 |
| **Marco** | | 1.00 |
| **Rin2** | | 1.00 |
| **Ccr2** | | 1.00 |
| **Coasy** | | 1.00 |
| **Tuba1a** | | 1.00 |
| **Tnfrsf8** | | 1.00 |
| **Acot11** | | 1.00 |
| **Sptbn1** | | 1.00 |
| **Itga4** | | 1.00 |
| **Nectin1** | | 0.83 |
| **Ccnb2** | | 1.00 |
| **Serpinb7** | | 1.00 |
| **Ptx3** | | 1.00 |
| **En1** | | 1.00 |
| **Pdcd1** | | 1.00 |
| **Il6** | | 1.00 |
| **Mif** | | 1.00 |
| **Cdh11** | | 1.00 |
| **Csf2ra** | | 1.00 |
| **Armc1** | | 1.00 |
| **Irf8** | | 1.00 |
| **Itgb1** | | 1.00 |
| **Daglb** | | 1.00 |

| | | |
|---|---|---|
| **Edn1** | | 1.00 |
| **Adgre5** | | 1.00 |
| **Lag3** | | 1.00 |
| **H2-Ab1** | | 1.00 |
| **Fgfr1** | | 1.00 |
| **Csf3r** | | 1.00 |
| **Adora2a** | | 1.00 |
| **Adam19** | | 1.00 |
| **Pde4a** | | 1.00 |
| **Cd40lg** | | 1.00 |
| **Rab20** | | 1.00 |
| **Ptgs1** | | 1.00 |
| **Map2k4** | | 1.00 |
| **Hspg2** | | 1.00 |
| **Rgs6** | | 1.00 |
| **Plaur** | | 1.00 |
| **Il17ra** | | 1.00 |
| **Il10** | | 1.00 |
| **C4a** | | 1.00 |
| **Sp4** | | 1.00 |
| **Fgf10** | | 1.00 |
| **Rgs16** | | 0.89 |
| **Adamts9** | | 1.00 |
| **Arhgef28** | | 1.00 |
| **Xcr1** | | 1.00 |
| **Sema4a** | | 1.00 |
| **Traf6** | | 1.00 |
| **Tnfrsf1b** | | 1.00 |
| **Cd44** | | 1.00 |
| **Bcl10** | | 1.00 |
| **Mertk** | | 1.00 |
| **Cxcr2** | | 0.83 |
| **Ifnar1** | | 1.00 |
| **Il5ra** | | 1.00 |
| **Isg15** | | 1.00 |
| **Serinc2** | | 1.00 |
| **Nox4** | | 1.00 |
| **Il22** | | 1.00 |
| **Pglyrp1** | | 1.00 |
| **Ido1** | | 1.00 |
| **Il1rn** | | 1.00 |
| **Rhoj** | | 1.00 |
| **Tlr4** | | 1.00 |
| **Kit** | | 1.00 |
| **Fcgr2b** | | 1.00 |
| **Tnfaip6** | | 1.00 |
| **Trem1** | | 1.00 |
| **Psmb9** | | 1.00 |

| | | |
|---|---|---|
| Cd209e | | 1.00 |
| Fadd | | 1.00 |
| Icosl | | 1.00 |
| Cyp1b1 | | 1.00 |
| Fzd4 | | 1.00 |
| Id3 | | 1.00 |
| Ptafr | | 1.00 |
| Fabp4 | | 1.00 |
| Ccr5 | | 1.00 |
| Mapk12 | | 1.00 |
| Hpgd | | 1.00 |
| 2900026A02Rik | | 1.00 |
| Ifngr1 | | 1.00 |
| Ltb4r1 | | 1.00 |
| Insr | | 1.00 |
| Cxcr3 | | 1.00 |
| Terf2ip | | 1.00 |
| Prok2 | | 1.00 |
| Mrc1 | | 1.00 |
| Klf4 | | 1.00 |
| Nox3 | | 1.00 |
| Nectin2 | | 1.00 |
| Hmgb1 | | 1.00 |
| Cd84 | | 1.00 |
| Stat3 | | 1.00 |
| Col3a1 | | 0.51 |
| Tyrobp | | 1.00 |
| Map2k2 | | 1.00 |
| Tlr13 | | 1.00 |
| Sphk1 | | 1.00 |
| Timp3 | | 1.00 |
| Fgf7 | | 1.00 |
| Cxcl5 | | 1.00 |
| Akap13 | | 1.00 |
| Ccl7 | | 1.00 |
| Ms4a2 | | 0.83 |
| Lipa | | 1.00 |
| Birc3 | | 1.00 |
| Itga5 | | 1.00 |
| Havcr2 | | 1.00 |
| Pbx1 | | 1.00 |
| Ccl1 | | 0.83 |
| Clec7a | | 1.00 |
| Il27 | | 1.00 |
| Trafd1 | | 1.00 |
| Kitl | | 1.00 |
| Gata6 | | 1.00 |
| Cpa3 | | 1.00 |

| | | |
|---|---|---|
| **Ltb4r2** | | 1.00 |
| **Fcer2a** | | 1.00 |
| **Mmp8** | | 1.00 |
| **S100a10** | | 1.00 |
| **Ldlr** | | 1.00 |
| **Tpsb2** | | 0.83 |
| **Mmp12** | | 1.00 |
| **Map2k1** | | 1.00 |
| **Tlr12** | | 1.00 |
| **Vcam1** | | 1.00 |
| **Myd88** | | 1.00 |
| **Nfkbie** | | 1.00 |
| **C1qb** | | 0.37 |
| **Elane** | | 1.00 |
| **Tnfsf10** | | 1.00 |
| **Sema5a** | | 1.00 |
| **Fpr1** | | 1.00 |
| **Ddr2** | | 1.00 |
| **Fn1** | | 1.00 |
| **Pycard** | | 1.00 |
| **Vasp** | | 1.00 |
| **Alox15** | | 1.00 |
| **Ifit1bl1** | | 1.00 |
| **Lta** | | 0.83 |
| **Tap2** | | 1.00 |
| **Nfyc** | | 1.00 |
| **Smarcd3** | | 1.00 |
| **Col10a1** | | 1.00 |
| **Trex1** | | 1.00 |
| **Peli1** | | 1.00 |
| **Gem** | | 1.00 |
| **Cd83** | | 1.00 |
| **Nkg7** | | 1.00 |
| **Ear6** | | 1.00 |
| **Cebpb** | | 1.00 |
| **Ikbke** | | 1.00 |
| **Kcnab1** | | 0.83 |
| **Col4a2** | | 1.00 |
| **Zfp92** | | 1.00 |
| **Ido2** | | 1.00 |
| **Ctgf** | | 1.00 |
| **Fcgr3** | | 0.83 |
| **Cdkn1a** | | 1.00 |
| **Abcc8** | | 1.00 |
| **Msc** | | 1.00 |
| **Pdgfra** | | 1.00 |
| **Il1r1** | | 1.00 |
| **Stat4** | | 1.00 |

| | | |
|---|---|---|
| **Socs2** | | 1.00 |
| **Scin** | | 1.00 |
| **Nfkbia** | | 1.00 |
| **Adamts14** | | 1.00 |
| **Siglecf** | | 1.00 |
| **Ctsl** | | 1.00 |
| **Insig1** | | 1.00 |
| **Ccl19** | | 1.00 |
| **Batf** | | 1.00 |
| **Prrx1** | | 1.00 |
| **Gtf2h2** | | 1.00 |
| **Dusp2** | | 1.00 |
| **Retnla** | | 1.00 |
| **Pggt1b** | | 1.00 |
| **Tigit** | | 0.37 |
| **Mpeg1** | | 1.00 |
| **Ccl12** | | 1.00 |
| **Rictor** | | 1.00 |
| **Was** | | 1.00 |
| **Sema6d** | | 1.00 |
| **Anxa1** | | 1.00 |
| **Nubp1** | Housekeeping | 1.00 |
| **Abcf1** | Housekeeping | 1.00 |
| **Ppia** | Housekeeping | 1.00 |
| **Sdha** | Housekeeping | 0.96 |
| **Rpl19** | Housekeeping | 1.00 |
| **Polr2a** | Housekeeping | 1.00 |
| **Oaz1** | Housekeeping | 1.00 |
| **Sf3a3** | Housekeeping | 1.00 |
| **Hdac3** | Housekeeping | 1.00 |
| **Alas1** | Housekeeping | 1.00 |
| **Tbp** | Housekeeping | 1.00 |
| **G6pdx** | Housekeeping | 1.00 |
| **Eef1g** | Housekeeping | 1.00 |
| **Tubb5** | Housekeeping | 1.00 |
| **Eif2b4** | Housekeeping | 1.00 |
| **Polr1b** | Housekeeping | 1.00 |
| **Edc3** | Housekeeping | 1.00 |
| **Hprt** | Housekeeping | 1.00 |
| **Sap130** | Housekeeping | 1.00 |
| **Gusb** | Housekeeping | 1.00 |
